# Thirty years of deforestation within the entire ranges of nine endangered lemur species (3 CR, 4 EN, 2 VU) in northwestern Madagascar

**DOI:** 10.1101/2022.05.23.493111

**Authors:** Dominik Schüßler, Yves Rostant Andriamalala, Robin van der Bach, Cheyenne Katzur, Christoph Kolbe, Mamy Hasina Rabe Maheritafika, Misa Rasolozaka, Mamy Razafitsalama, Marten Renz, Travis S. Steffens, Ute Radespiel, Jana Brenner

## Abstract

Forest cover change is of particular concern in tropical regions. In this study, we investigate the degree of deforestation in the entire ranges of nine highly threatened lemur species in northwestern Madagascar. Landsat satellite images were acquired from four different time stages (1990, 2000, 2011, 2020), classified into forest/non-forest, and changes quantified. Forest cover declined from 17.5% to 9.3% within the last 30 years. This decline varied across four protected areas (PAs) investigated: the forest cover of Ankarafantsika National Park (ANP) declined only moderately over time (from 76.3% to 67.4%), while it declined drastically in other PAs (e.g., from 54.9% to 18.9%, Bongolava Forest Corridor). Two lemur taxa are most affected (*Lepilemur otto, Microcebus bongolavensis*) by having only very few isolated forest patches left within their ranges (approximately 542.7 km²). For two other species (*L. ahmansoni, L. aeeclis*), most of the remaining forest is concentrated in two coastal PAs (in total 627.2 and 477.9 km², respectively), while those species occurring inside ANP (5 taxa) experienced rather stable forest coverage until 2020. A reversal of these deforestation trends and active reforestation measures are desperately needed to reduce habitat loss for these nine lemur species. A practical experience-based guideline is therefore provided.

## Introduction

Forest cover patterns in Madagascar over time have long been disputed. However, understanding these patterns and the processes that created them is pivotal for understanding today’s species distributions, past extinctions and the anthropogenic role in this process. Contrary to a long held hypothesis of an entirely forest-covered Madagascar prior to human arrival (e.g., Gade 1996, see Kull 2000), evidence is accumulating that most parts of Madagascar, especially the central highlands, were more likely to have been characterized by a grass-woodland mosaic, with more or less forest cover depending on the paleoclimatic conditions and the amount of precipitation (Crowley et al. 2021, Joseph et al. 2020, Solofondranohatra et al. 2018). Following this hypothesis, the increasing aridity (except in the eastern rainforests) since the end of the African Humid Period at around 5.5k years before present (DeMenocal et al. 2000) was one of the major drivers for the late Holocene decline in forest cover and subsequent demographic changes for a variety of species (Gasse & van Campo 2001, Teixeira et al. 2021a,b, Salmona et al. 2017, Quéméré et al. 2012). These natural processes were accompanied with habitat fragmentation effects triggered by an expanding human population and a shift of livelihood activities towards cattle herding and agriculture within the last 1-2k years (Godfrey et al. 2019, Vorontsova et al. 2016).

Since the middle of the 20^th^ century, it is possible to monitor changes in forest cover consistently across the island by means of remote sensing data. Starting with aerial photographs from the 1950s (Harper et al. 2007, Scales 2011), and continued with Landsat satellite images from the 1970s onwards (Vieilledent et al. 2018), the degree of mainly human induced forest cover change in recent times can be assessed. Deforestation between 1973 and 2014 contributed to a decrease in forest cover in the dry deciduous forests of western Madagascar of 41.5% (4,435 to 2,596 kha) with the period since 1990 accounting for almost half of this decline (19.5% decline 1990-2014). This has led to increasing fragmentation as indicated by decreasing mean distances to forest edges (1.5km in 1973 to 300m in 2014) and about 46% of core forest being located less than 100m away from the edge (Vieilledent et al. 2018). This has unmistakable consequences for people and wildlife depending on forest-derived ecosystem services. Consistent with paleontological records and the assembly of extant species, most wildlife in Madagascar was and is adapted to a forest-dwelling lifestyle, requiring a certain degree of tree coverage (Goodman & Benstead 2005). The lemurs of Madagascar are a striking example for the consequences of forest cover change (i.e., loss) with 98% of all known 100+ species being listed as threatened with extinction by the IUCN (www.iucnredlist.org). Negative effects of forest fragmentation have been documented in a variety of ecological and genetic studies for different lemur species (e.g., Andriatsitohaina et al. 2019, Craul et al. 2009, Salmona et al. 2017, Quéméré et al. 2012).

In this study, we provide a detailed analysis of the forest cover changes within the entire ranges of nine threatened lemur species in northwestern Madagascar over the last 30 years to highlight the urgent need to effectively protect certain key habitats from further deforestation in order to prevent the extinction of these and other co-occurring species. We end our study with a practical guide on lemur-focused forest restoration.

## Methods

### Study region

The study region has a size of 69,290.35 km² and was defined by the last remaining larger forest complex of northwestern Madagascar (Ankarafantsika National Park, ANP) and three adjacent inter-river systems covered by cloud-free areas of two neighboring Landsat satellite images available for four different time intervals (1990-2020, Fig. 1). We thereby covered the entire ranges of nine partly sympatric, forest-dwelling lemur species (Table 1). A further ten species also occur in this region, with their ranges extending beyond it but are nevertheless negatively impacted by forest cover loss (i.e., *Cheriogaleus medius, Daubemtonia madagascariensis, Eulemur fulvus, E. rufus, Hapalemur griseus, Lepilemur grewcockorum, Microcebus danfossi, M. murinus, M. myoxinus, Propithecus coronatus*). Four protected areas are located in the region. These are Antrema NPA (new protected area), Ankarafanstika NP (National Park), Bongolava FC (forest corridor) and the Mahavavy-Kinkony wetlands NPA. The region is primarily characterized by dry deciduous forests, open grassland vegetation and agricultural activities (Gade 1996, Koechlin 1972). Several large rivers with their headwaters in the central highlands traverse northwestern Madagascar allowing to define different inter-river systems (Fig. 1). These major rivers are considered as distributional boundaries for a variety of species (e.g., Craul et al. 2007, Olivieri et al. 2007).

**Table 1:** Lemur species with their entire ranges within the study region and their IUCN conservation states (Mittermeier et al. 2010, www.iucnredlist.org). IRS = Inter-river system as defined by Craul et al. (2007), see Figure 1, CR = Critically Endangered, EN = Endangered, VU = Vulnerable.

| Lemur species |  | IUCN | Distribution in |
| --- | --- | --- | --- |
|  |  | status | IRS |
| *Avahi occidentalis* | Western wooly lemur | VU | I, II |
| *Eulemur mongoz* | Mongoose lemur | CR | –I, 0, I, II |
| *Lepilemur aeeclis* | Antafia sportive lemur | EN | 0 |
| *Lepilemur ahmansoni* | Tsiombikibo sportive lemur | CR | –I |
| *Lepilemur edwardsi* | Milne-Edwards sportive lemur | EN | I |
| *Lepilemur otto* | Ambodimahadibo sportive lemur | EN | II |
| *Microcebus bongolavensis* | Bongolava mouse lemur | EN | II |
| *Microcebus ravelobensis* | Golden-brown mouse lemur | VU | I |
| *Propithecus coquereli* | Coquerel’s sifaka | CR | I-III |

**Figure 1:**
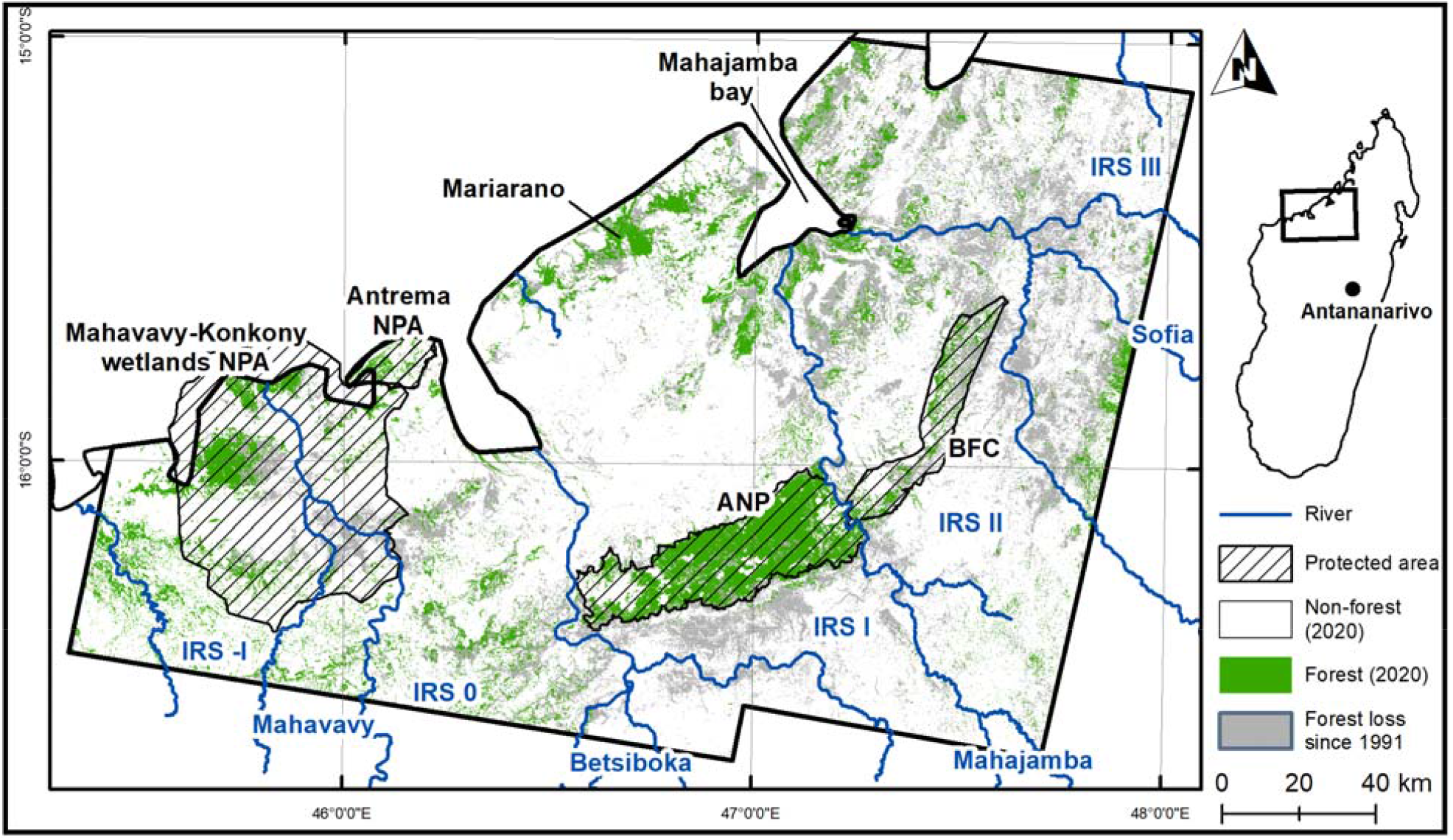
Forest cover in 2020 and forest loss since 1991 in northwestern Madagascar. Protected areas and inter-river systems (IRS) are marked for orientation. ANP = Ankarafantsika National Park, BFC = Bongolava Forest Corridor, NPA = New Protected Area.

### Forest cover mapping

Georeferenced and terrain corrected satellite images were retrieved from the U.S. Geological Survey (www.earthexplorer.usgs.gov) for four different time stages: 1990/91, 2000, 2011 and 2020 (Table S1). No radiometric or solar correction was applied, as satellite images were analyzed separately (i.e., we did not create mosaic images) and no derived indices were calculated. Different optical bands were combined to yield raster stacks of 30 m x 30 m spatial resolution (Table S1). We classified the satellite images applying a supervised maximum likelihood algorithm based on sets of training areas for each land cover class (Table S2). These were *a priori* defined as forest (= dense and broad-leafed vegetation with a nearly closed canopy), open soil (= almost barren land with only marginal grass coverage including dry agricultural fields (due to season)), grassland (= dense grass coverage with interspersed trees) and water (= open water bodies like lakes, rivers and the Indian Ocean). Land cover classes were *a posteriori* summarized as forest and non-forest to increase the accuracy of the classification. Areas for training of the algorithm were selected based on our own experiences from the region and from published records (Andriatsitohaina et al. 2019, Rakotondravony & Radespiel 2009). Annual rate of forest cover change (i.e., mean annual deforestation or afforestation) was calculated following formula (7) of Puyravaud (2003).

Accuracy assessment was conducted using 500 random points across the study region in a stratified sampling scheme (i.e., proportional number of points according to land cover class) for each time interval. Ground truthing was based on visual interpretation of actual land cover (forest or non-forest) as displayed in the Landsat images at each random point based on experiences from previous projects (Schüßler et al. 2018, 2020). User’s and producer’s accuracy (commission and omission rate, respectively) and the overall accuracy were calculated according to guidelines by Olofsson et al. (2014). All analyses were conducted in ArcMap (ArcGIS Desktop 10.6.1, ESRI, Redlands, USA). The forest cover maps for the different time stages are freely available and can be downloaded at: https://doi.org/10.25625/2U5U5J

## Results

Overall accuracies for the four time stages 1990/91, 2000, 2011 and 2020 were 92.2%, 91.4%, 93.0% and 96.0%, respectively. User’s and Producer’s accuracies did not fall below 71.4% and 83.3% across all time stages, respectively (Table S3).

### Forest cover change

The overall trend of forest cover change was negative, with a decline from 17.52% in 1991 to 9.26% in 2020. This was, however, inverted between 2000 and 2011 with a slight increase in forest cover (Table 2). This observation can be exemplified in areas at the northern border of ANP, where regenerating vegetation in 2000 was not classified as forest, but gained comparable reflectance values as forests by 2011 (Fig. 2, Fig S1). Mean annual rates of forest change were 2.54% deforestation (1991-2000), 1.11% afforestation (2000-2011) and 1.89% deforestation (2011-2020). Within the four protected areas, trends were different: In Ankarafantsika NP, it was fluctuating over time, with regenerating forests until 2011 and a decline towards 2020 (Fig. 2). In Antrema NPA, we observed an increase in forest cover since 1991, while in Bongolava FC, we documented the strongest and most pronounced deforestation from >54% in 1991 to <19% forest coverage in 2020. In Mahavavy-Kinkony wetlands NPA instead, forest cover was rather stable but declined sharply towards 2020 (Table 2). This primarily happened in the southeastern part of the protected area, where riparian forests were converted into agricultural fields (Fig. 2).

**Table 2:** Forest cover in km² (and percentages in brackets) in the study region and inside the four protected areas between 1991-2020.

|  | Size [km<sup>2</sup>] | 1991 | 2000 | 2011 | 2020 | Protected since |
| --- | --- | --- | --- | --- | --- | --- |
| Study region | 69,290.35 | 12,141.19<br>(17.5%) | 7,169.66<br>(10.4%) | 9,485.31<br>(13.7%) | 6,417.95<br>(9.3%) | - |
| Ankarafantsika NP | 1,351.24 | 1,031.59<br>(76.3%) | 914.88<br>(67.7%) | 1,020.91<br>(75.6%) | 911.34<br>(67.4%) | 1927 |
| Antrema NPA | 190.46 | 29.34<br>(15.4%) | 35.61<br>(18.7%) | 45.70<br>(24.0%) | 43.91<br>(23.1%) | 2010 |
| Bongolava FC | 607.00 | 333.13<br>(54.9%) | 258.63<br>(42.6%) | 186.63<br>(30.7%) | 114.40<br>(18.9%) | 2015 |
| Mahavavy-Kinkony wetlands NPA | 2,995.84 | 651.99<br>(21.8%) | 874.19<br>(29.1%) | 873.54<br>(29.1%) | 427.87<br>(14.3%) | 2007 |

**Figure 2:**
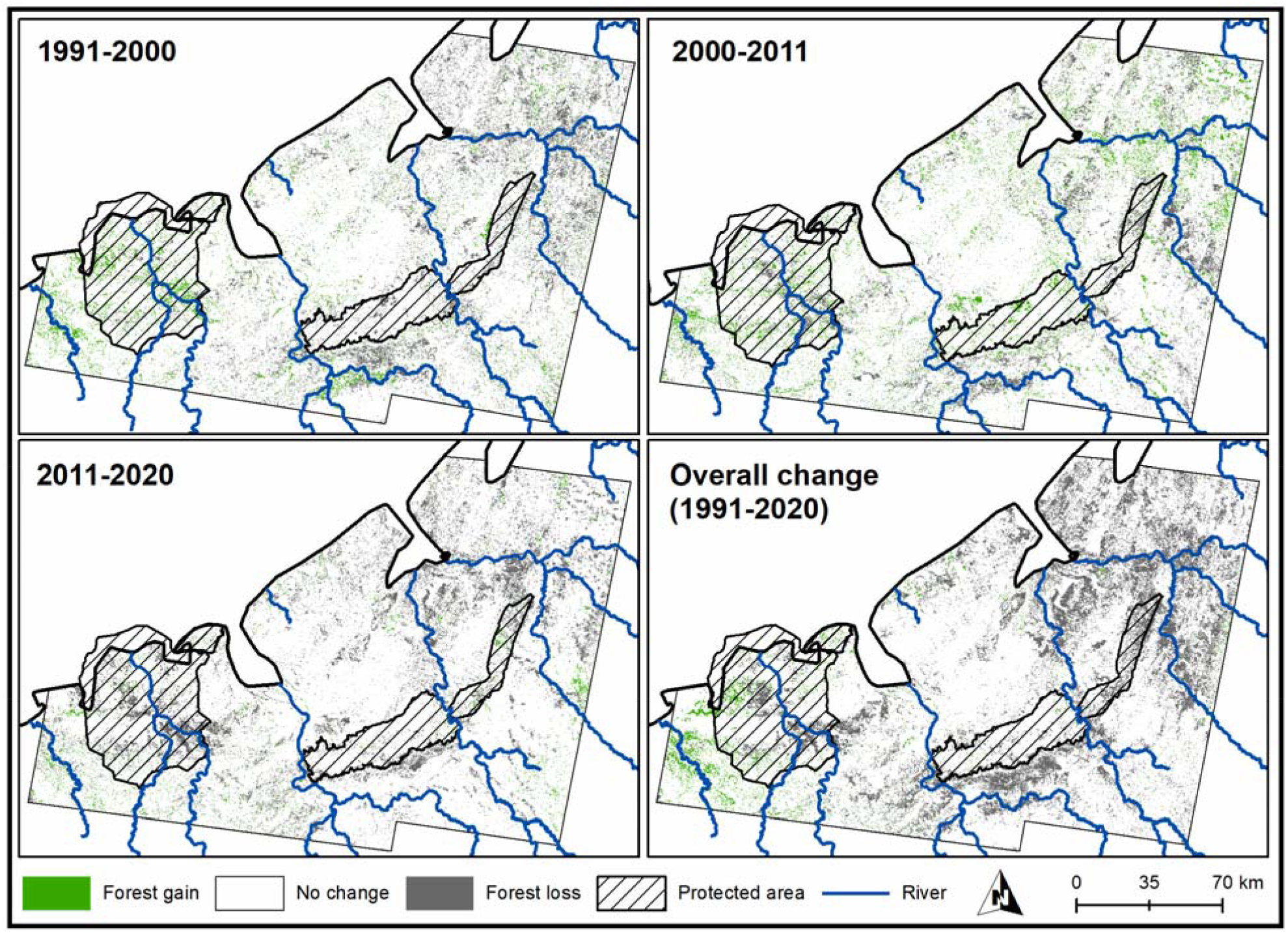
Forest gain and loss from 1991-2020. Protected area names and labels of inter-river systems are given in Figure 1.

### Impact on lemur populations

Concerning the ranges of the nine lemur species, those taxa occurring in IRS I (*A. occidentalis, E. mongoz, L. edwardsi, M. ravelobensis, P. coquereli*) have the highest proportion of forested habitats still available within their ranges (1,535.2 km² in IRS I). These are located in Ankarafantsika NP and along the coast around the village of Mariarano (Fig. 1). Apart from smaller forest patches on the southwestern stretch of the Mahajamba bay, these two forested areas have become increasingly disconnected from each other since 1991. For species with their entire distribution confined to IRS II (*L. otto, M. bongolavensis*), only very few small forest fragments remain in 2020 totaling with a size of 552.7 km² (6.3% of IRS II, Table 3). These are partly located inside Bongolava FC where forest cover decreased by 36% since 1991 (Table 2) and along the Mahajamba bay (Fig. 1). IRS 0 and hence the range of *L. aeeclis* harbors the last remaining forests with a total size of 477.9 km² (9.3% of IRS 0) along the coast within the two protected areas and in a few unprotected stretches further inland. This also applies to IRS -I and the range of *L. ahmansoni* with a total amount of forest cover of 627.2 km² (15.5% of IRS –I, Table 3). However, further small forest patches can be found west of Mahavavy-Kinkony wetlands NPA (Fig 1).

**Table 3:** Forest cover change in km² (percentages in brackets) in the different inter-river systems (IRS) in relation to the ranges of the concerned lemur species. Species with their entire range confined to a single IRS are given in bold.

| IRS | Size | 1990/91 | 2000 | 2011 | 2020 | Species concerned |
| --- | --- | --- | --- | --- | --- | --- |
| -I | 4,036.29 | 607.67<br>(15.1%) | 895.58<br>(22.2%) | 961.03<br>(23.8%) | 627.18<br>(15.5%) | *E. mongoz*, ***L. ahmansoni*** |
| 0 | 5,166.39 | 1093.10<br>(21.2%) | 1050.99<br>(20.3%) | 947.38<br>(18.3%) | 477.88<br>(9.3%) | *E. mongoz*, ***L. aeeclis*** |
| I | 11,064.88 | 2992.49<br>(27.0%) | 2442.37<br>(22.1%) | 2434.02<br>(22.0%) | 1535.22<br>(13.9%) | *E. mongoz*, *A. occidentalis*, ***L. edwardsi***, ***M. ravelobensis***, *P. coquereli* |
| II | 8,685.78 | 2715.74<br>(31.3%) | 1849.17<br>(21.3%) | 1477.11<br>(17.0%) | 542.72<br>(6.3%) | *E. mongoz*, *A. occidentalis*, ***L. otto***, ***M. bongolavensis***, *P. coquereli* |

## Discussion

### Forest cover change

Deforestation was an ongoing trend in Madagascar for many decades (Vieilledent et al. 2018) and declines in the dry deciduous forests of western Madagascar were already considered to require an substantial increase in attention (Waeber et al. 2015). Compared to the whole ecological region, mean annual deforestation rates were markedly higher and comparable in the first and third study decade (1991-2000, 2011-2020) within the study region. Between 2000-2011, however, a net-recovery of forest cover was documented, a trend not recorded by a larger scale dataset (i.e., Vieilledent et al. 2018) or observations from the Menabe region of western Madagascar (Scales 2011). For example, a significant recovery of forest was observed at the northern border of Ankarafantsika NP, where a larger section of forest was lost by 2000 but appeared to have recovered by 2011. As detailed in supplementary figure S1, we attribute this loss to a wildfire which must have occurred in that area before 2000. Visual interpretation of the false-color composites of the satellite images indicated natural rejuvenation of previously burned places in comparison with surrounding areas. Apart from this region, significant forest recovery could be observed and verified from raw satellite images in the Tsiombikibo forest in northwestern section of Mahavavy-Kinkony wetlands NPA and in northern parts of IRS II. We therefore conclude that the observed forest recovery was not an analytical artifact due to a slightly earlier sensing date of the Landsat images in 2011 (July instead of August/September 2000), but rather related to the process of forest recovery following fire.

Forest cover change inside the four protected areas followed different trends, only partly reflected by the changes in the overall study region. For Bongolava FC, we documented the strongest and steadiest decline. Ankarafantsika NP fluctuated around a high proportion of forest within its boundaries, while in Antrema NPA forest cover increased and stabilized over time. In Mahavavy- Kinkony wetlands NPA, forests first recovered in one section before a strong decline was observed in the south of this protected area in favor of agricultural expansion. Unfortunately, this area was one of those with highest densities of Critically Endangered *Propithecus coronatus* (Salmona et al. 2014). Differential forest cover change in various protected areas are also mirrored by other studies from Madagascar (Eklund et al. 2016, Rafanoharana et al. 2021) highlighting the region-depended challenges and differences in protected area management.

Although historic forest cover change in Madagascar was most likely triggered by climatic changes (Crowley et al. 2021, Joseph et al. 2020, Solofondranohatra et al. 2018), recent deforestation in the study region is mainly driven by human subsistence and partly commercial activities like agricultural expansion, livestock grazing or forest exploitation for charcoal production (Dave et al. 2016, Gay-des-Combes et al. 2017, Jones et al. 2015, Steffens et al. 2020, this study). While being in accordance with other regions of sub-Saharan Africa (Curtis et al. 2018), local land users in northwestern Madagascar and elsewhere are highly dependent on particular forest-related ecosystem services to sustain their livelihoods, but increasingly notice the decline of these services (Dave et al. 2016, Steffens et al. 2020). Our analysis provides remotely sensed evidence to the experiences of residents from the region: forest cover and associated ecosystem services (i.e, the provision of raw materials or storm hazard mitigation) has been steadily declining for decades now (Dave et al. 2016, Steffens et al. 2020), with no indications for a trend reversal.

In fact, after the acquisition date of the latest satellite image used for our analysis (September 2020), and during the finalization of this article in December 2021, Ankarafantsika NP was hit hard by extensive fires. Forest loss equaled 6.66% of the park’s surface or roughly 90 km² - within one year of time (Fig. 3). It is to be hoped, that natural regeneration processes can be initiated and land degradation towards grasslands can be counteracted, to safeguard forest connectivity inside this key protected area of northwestern Madagascar.

**Figure 3:**
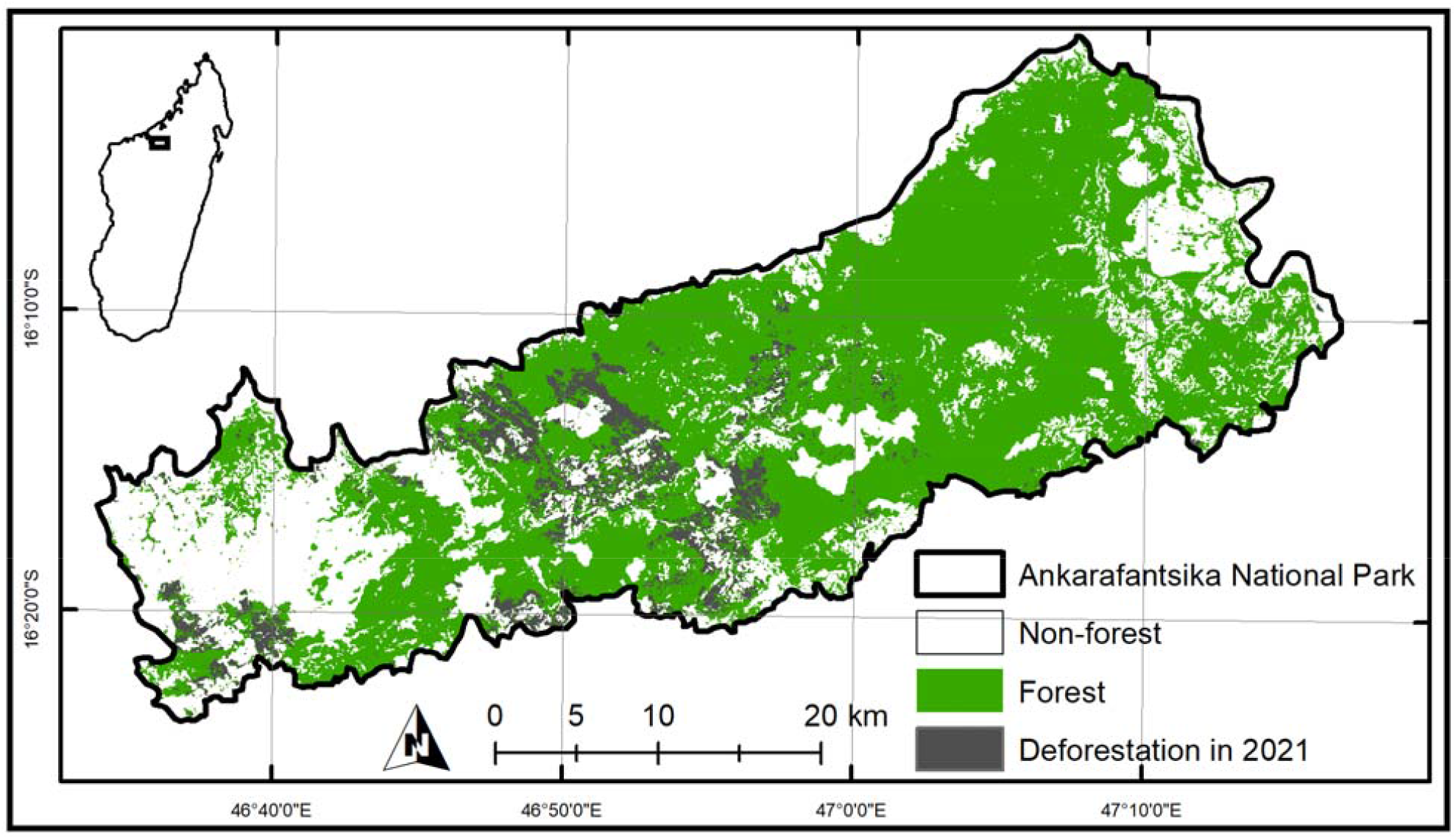
Forest burning inside Ankarafantsika NP between September 2020 and December 2021 as derived from Sentinel-2 satellite images acquired on 04-12-2021 (tiles: T38-LQH, T38-KPH).

### Impact on lemur populations

For the majority of lemur species, the population-wide influence of habitat loss and fragmentation is not well investigated yet (Hending 2021, Kling et al. 2020). For those species having parts of their distributions inside Ankarafantsika NP (in IRS I), the proportion of available habitat was relatively stable in its extent until 2021. The impacts of the latest extensive fires have to be examined in the future. The previous stability in forest coverage was mirrored in the highest degree of genetic diversity for *P. coquereli* when compared to other species from this genus (Bailey et al. 2016). However, even in this rather stable region of northwestern Madagascar, local community members reported decreasing lemur populations and the diminishment of natural resources during their lifetimes (Steffens et al. 2020). The situation for species outside of IRS I is much more severe. Forest cover in the ranges of *L. aeeclis, L. ahmansoni, L. otto* and *M. bongolavensis* declined drastically leaving only very few small refuges for these species today. It has to be noted, that our figures of extent forest cover can only represent the maximum size of available habitats, leaving out habitat quality and the patchiness of the remaining forests.

Several studies on fragmentation effects give insights into the possible underlying patterns that can be linked to forest cover changes as inferred by our analysis (e.g., Eppley et al. 2020). The lemur species of northwestern Madagascar are in general not equally distributed across forest fragments with notable absences mainly of the larger bodied species or folivorous and arboreal nocturnal taxa (*Avahi* spp., *E. mongoz, Lepilemur* spp.) from certain fragments (Olivieri et al. 2005). The size of a particular forest was found to be more important than fragment isolation in determining the species richness of lemurs (Steffens & Lehmann 2018, 2019). For example, *P. coquereli* and *L. edwardsi* did not occur in smaller fragments and did not form metapopulations across fragments (Steffens & Lehman 2018) while *A. occidentalis* and *E. mongoz* were not found in fragments around but in Ankarafantsika NP itself (Steffens & Lehman 2018, Steffens et al. 2020). Apart from this classical island biogeographic perspective (e.g., Diamond 1985), several species are negatively impacted by forest fragmentation, displayed by lower genetic diversity or recent population collapses (*L. edwardsi*, Craul et al. 2009, *M. bongolavensis* and *M. ravelobensis*, Olivieri et al. 2008, *M. ravelobensis*, Radespiel et al. 2018, Andriatsitohaina et al. 2019, *P. coquereli*, Bailey et al. 2016) and possibly by molecular edge effects (*M. ravelobensis*, Radespiel et al. 2018). Furthermore, whereas endemic species appear to be negatively impacted, non-native species like *Rattus rattus* benefit from increasing fragmentation (Andriatsitohaina et al. 2019). If the described trends cannot be reversed, local extirpations and genetic impoverishment have to be expected.

### Lemur-focused forest restoration – a practical guide

In order to prevent population extinctions and to invert the ongoing loss of forest cover, restoration efforts are needed to maintain the ecosystem services people and wildlife rely on.

The non-governmental organization Planet Madagascar and it’s implementing partner Planet Madagascar Association have been restoring forests around Ankarafantsika NP since 2017. Their main goal is to protect and to restore forest corridors for lemurs while providing opportunities for sustainable livelihoods for people that share these forests. Forest restoration can be effectively done following their experience-based 6-step framework:

1. Plantation site selection is key to ensure restoration success. Planet Madagascar combined local knowledge with scientific data on the occupancy of lemurs, major threats to the forests within their management zone, and logistical considerations (e.g., proximity to communities and water) to determine which portion of the landscape should be restored. The restoration site was selected because (a) it contained forest fragments and adjacent continuous forest with high lemur species diversity (Steffens and Lehman 2018, 2019, Andriatsitohaina et al. 2019) and (b) because forests fragments were in close proximity (i.e., ∼ 500 m) to allow reconnection via forest re-growth. Additionally, it is important (c) to incorporate advantageous landscape features like natural depressions (i.e., low lying areas) where water retention capacity is naturally increased.
2. Species selection is integral to ensure recruitment. Plant species must be selected that are used by lemurs and that are abundant in nearby continuous forest (Supplementary Table S4). The selected species should grow relatively fast and be native to the area to ensure adaptability to the local climate conditions or non-invasive (e.g. *Tamarindus indica*).
3. Nursing of seedlings increases recruitment. This step involves three parts: (a) seed collection, (b) seed processing, and (c) nursing conditions.
  a. Fresh seeds must be collected from June to December (dry season) and planted immediately into a nursery. Storing of seeds is not advised as they can be preyed upon by insects or destroyed by mold. In 2017-2018 for example, Planet Madagascar lost more than 75% of seedlings that were stored from October to February.
  b. Recruitment of seedlings can be increased when seeds are rinsed prior to planting. For example, Planet Madagascar compared recruitment of *Rhopalocarpus similis* and *Vitex* spp. seedlings that were treated by rinsing only and rinsing plus soaking in water for 24 hours. Rinsed seeds of both species had 31% and 95% higher recruitment than rinsed and soaked seeds, respectively.
  c. Nurseries must be placed in areas with high quality soils and adjacent to permanent water supply. Poor soil quality has negative impacts on recruitment and quality of saplings. Nurseries on good soils produced 4% more saplings with all being of bigger size and girth.
4. Planting of saplings from the nurseries to receiver areas must be done during the rainy season (December to late March in ANP).
5. Preparation of the plantation improves sapling survivorship. Planet Madagascar usually prepares the plantation area by clearing a firebreak around it, digging small holes for the saplings, clearing grass around each hole, and adding compost. Planting in proximity to shade trees or installation of artificial shade providers improves sapling survivorship. For example, saplings under shade had more than 70% greater survivorship than without shade. Additional measure to protect against damage from cattle (i.e. fencing) are recommended if financially feasible.
6. Continued monitoring of planted saplings is required to counteract unexpected events and prevent cattle from grazing on saplings. Fire is a common cause of failure for forest restoration projects (Elliott et al. 2013). Clearing plantations from encroaching grass during monitoring is therefore considered important to maintain the fire breaks and to reduce competition for nutrients between grasses and saplings.

Within a duration of three years (December 2017-2020), Planet Madagascar has planted 66,937 seedlings into a plantation with a survivorship of 31.46% (21,061 seedlings) restoring 100 ha of dry deciduous forest around Ankarafantsika NP. Future research should further assess other ways of improving reforestation methods, long-term survivorship of trees, and whether lemurs actually use these restored forest as habitat.

## Acknowledgements

This project did not receive external funding and was not directly linked to fieldwork activities in Madagascar. We warmly thank the village community members of Andranohobaka, Maevatanimbary, and Ambarindahy for engaging in the forest restoration projects around Ankarafantsika National Park. Funds to support Planet Madagascar’s forest restoration initiatives were provided by the IUCN Save Our Species fund and through the Department of Foreign Affairs, Trade, and Development, Canada Fund for Local Initiatives.

## Supplementary Material

**Supplementary Table S1:**
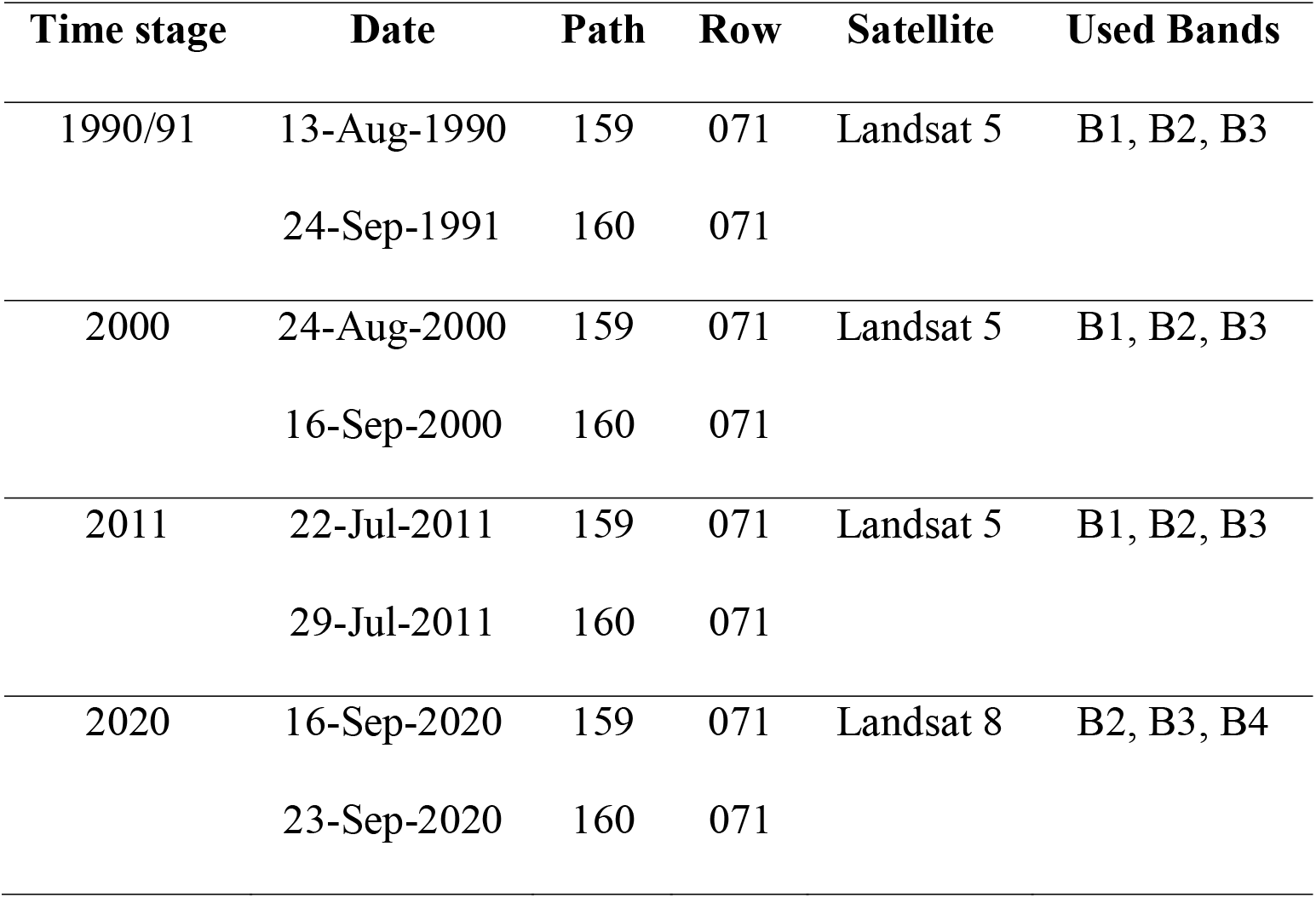
Technical details (sensing date and geographic location) of Landsat satellite images acquired from the U.S. Geological Survey.

| Time stage | Date | Path | Row | Satellite | Used Bands |
| --- | --- | --- | --- | --- | --- |
| 1990/91 | 13-Aug-1990 | 159 | 071 | Landsat 5 | B1, B2, B3 |
|  | 24-Sep-1991 | 160 | 071 |  |  |
| 2000 | 24-Aug-2000 | 159 | 071 | Landsat 5 | B1, B2, B3 |
|  | 16-Sep-2000 | 160 | 071 |  |  |
| 2011 | 22-Jul-2011 | 159 | 071 | Landsat 5 | B1, B2, B3 |
|  | 29-Jul-2011 | 160 | 071 |  |  |
| 2020 | 16-Sep-2020 | 159 | 071 | Landsat 8 | B2, B3, B4 |
|  | 23-Sep-2020 | 160 | 071 |  |  |

**Supplementary Table S2:** Number of training samples per class

| Time stage | Training samples | Forest | Disorders | Grassland | Ocean | Water | Erosion, rocks |
| --- | --- | --- | --- | --- | --- | --- | --- |
| 1990/91 | 44 | 20 | - | 6 | 3 | 8 | 7 |
| 2000 | 36 | 15 | - | 3 | 3 | 8 | 7 |
| 2011 East | 206 | 38 | - | 59 | 6 | 39 | 64 |
| 2011 West | 388 | 46 | 73 (clouds)<br>71 (shadow) | 56 | 5 | 70 | 67 |
| 2020 East | 158 | 39 | 42 (clouds) | 21 | 2 | 36 | 18 |
| 2020 West | 158 | 43 | - | 30 | 6 | 40 | 39 |

**Supplementary Table S3:** User’s, Producer’s and overall accuracies for the four different time stages per land cover class. Calculations based on 500 assessment points under a stratified random sampling (Olofsson et al. 2014).

| Time stage | Land cover class | Producer's accuracy | User's accuracy | Overall accuracy |
| --- | --- | --- | --- | --- |
| 1990/91 | Non-Forest | 0.932 | 0.965 | 0.922 |
|  | Forest | 0.891 | 0.803 |  |
| 2000 | Non-Forest | 0.933 | 0.960 | 0.914 |
|  | Forest | 0.833 | 0.748 |  |
| 2011 | Non-Forest | 0.946 | 0.968 | 0.930 |
|  | Forest | 0.857 | 0.780 |  |
| 2020 | Non-Forest | 0.967 | 0.989 | 0.960 |
|  | Forest | 0.884 | 0.717 |  |

**Supplementary Table S4:**
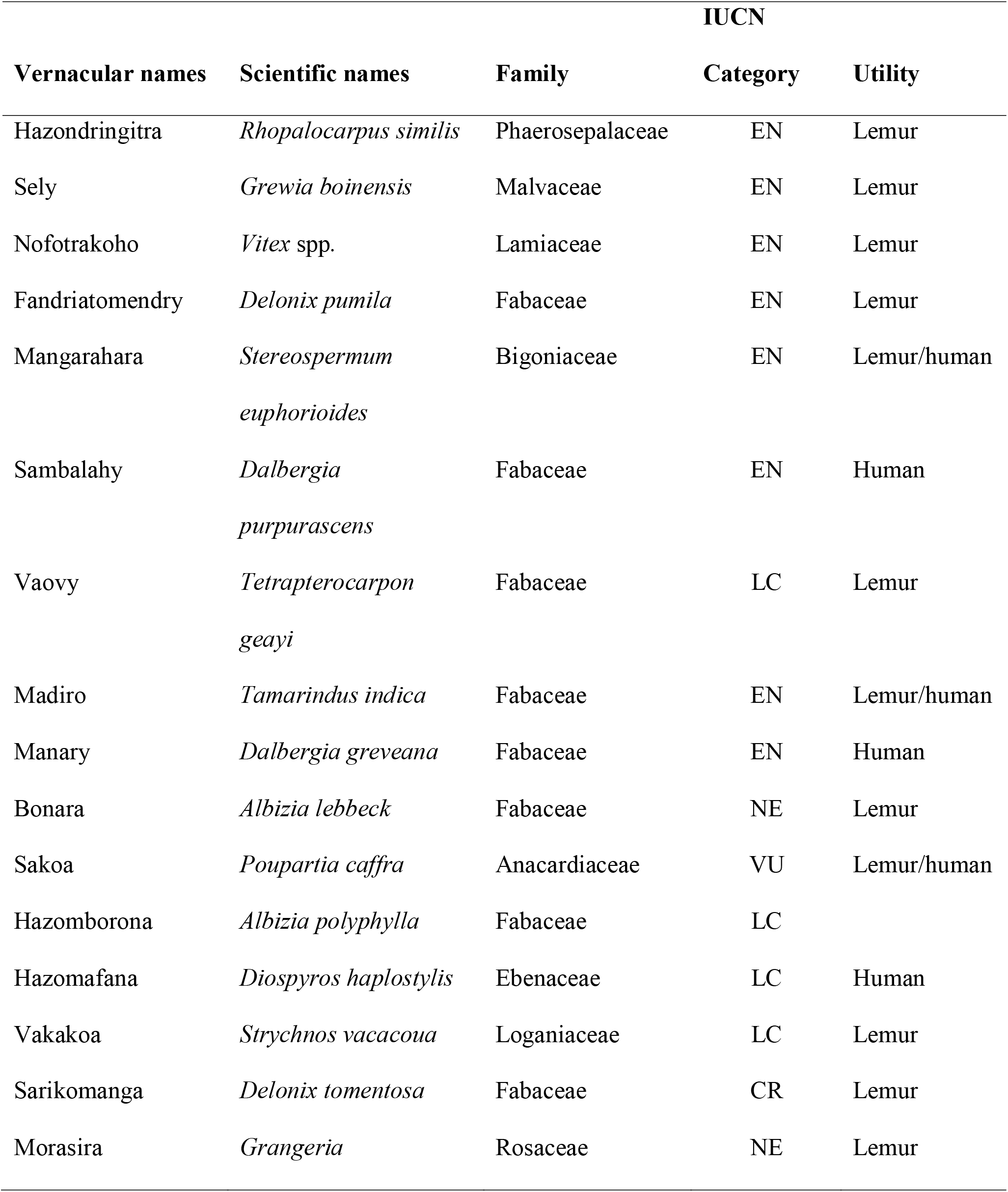
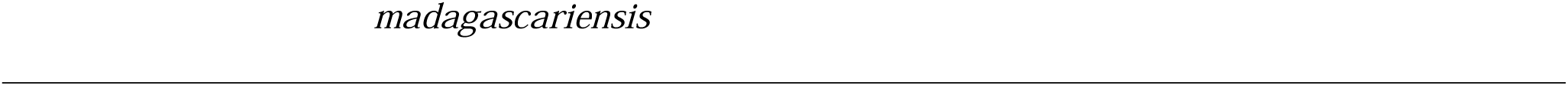
Tree species used for restoration with conservation states. LC = Least Concern, EN = Endangered, CR = Critically Endangered, NE = Not Evaluated.

| Vernacular names | Scientific names | Family | IUCN |  |
| --- | --- | --- | --- | --- |
|  |  |  | Category | Utility |
| Hazondringitra | *Rhopalocarpus similis* | Phaerosepalaceae | EN | Lemur |
| Sely | *Grewia boinensis* | Malvaceae | EN | Lemur |
| Nofotrakoho | *Vitex* spp. | Lamiaceae | EN | Lemur |
| Fandriatomendry | *Delonix pumila* | Fabaceae | EN | Lemur |
| Mangarahara | *Stereospermum*<br>*euphorioides* | Bigoniaceae | EN | Lemur/human |
| Sambalahy | *Dalbergia*<br>*purpurascens* | Fabaceae | EN | Human |
| Vaovy | *Tetrapterocarpon*<br>*geayi* | Fabaceae | LC | Lemur |
| Madiro | *Tamarindus indica* | Fabaceae | EN | Lemur/human |
| Manary | *Dalbergia greveana* | Fabaceae | EN | Human |
| Bonara | *Albizia lebbek* | Fabaceae | NE | Lemur |
| Sakoa | *Poupartia caffra* | Anacardiaceae | VU | Lemur/human |
| Hazomborona | *Albizia polyphylla* | Fabaceae | LC |  |
| Hazomafana | *Diospyros haplostylis* | Ebenaceae | LC | Human |
| Vakakoa | *Strychnos vacacoua* | Loganiaceae | LC | Lemur |
| Sarikomanga | *Delonix tomentosa* | Fabaceae | CR | Lemur |
| Morasira | *Grangeria* | Rosaceae | NE | Lemur |

**Supplementary Figure S1:**
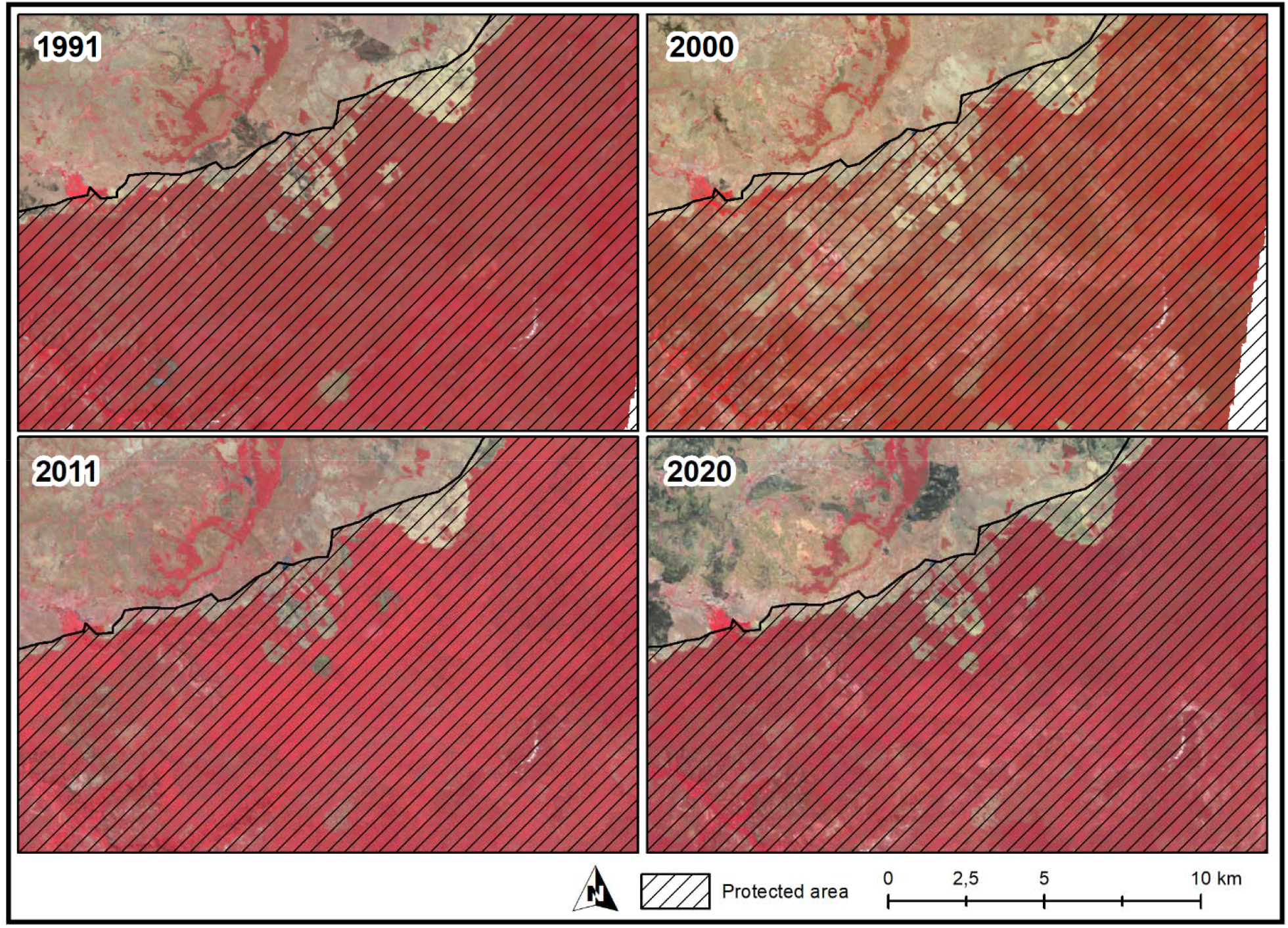
Detailed illustration of the appearance of satellite images (in false color) at the northern central section of Ankarafantsika National Park. Strong red colors indicate forest cover, beige color indicates open soil and black patches are recently burned places. Forest cover in 2000 was markedly reduced in this section but recovered to its former extent by 2011. Forest cover loss as it can be seen in the 2000 time stage was most likely due to a wildfire. In 2011, previously burned areas appear in stronger red color than unburned forests, indicating strong regrowth and stimulating effects of fire on the natural rejuvenation.

## Notes

https://doi.org/10.25625/2U5U5J

### Competing Interest Statement

The authors have declared no competing interest.

